# Quantifying Coral Morphology

**DOI:** 10.1101/553453

**Authors:** Kyle J. A. Zawada, Maria Dornelas, Joshua S. Madin

**Author notes:** Corresponding author: Kyle Zawada, Centre for Biological Diversity, Scottish Oceans Institute, University of St. Andrews, KY16 9TH, Scotland, UK.

## Abstract

The morphology of coral colonies has important implications for their biological and ecological performance, including their role as ecosystem engineers. However, given that morphology is difficult to quantify across many taxa, morphological variation is typically shoehorned into coarse growth form categories (e.g., arborescent and digitate). In this study, we develop a quantitative schema for morphology by identifying three-dimensional shape variables that can describe coral morphology. We contrast six variables estimated from 152 laser scans of coral colonies that ranged across seven growth form categories and three orders of magnitude of size. We found that 88% of the variation in shape was captured by two axes of variation and three shape variables. The main axis was variation in volume compactness (*cf*. sphericity) and the second axis was the trade-off between surface complexity and the vertical distribution of volume (i.e., top heaviness). Variation in volume compactness also limited variation along the second axis, where surface complexity and vertical volume distribution ranged more freely when compactness was low. Traditional growth form categories occupied distinct regions within this morpho-space. However, these regions overlapped due to shape changes with colony size. Nonetheless, four of the shape variables were able to predict traditional growth form categories with 70 to 95% accuracy, suggesting that the continuous variables captured much of the qualitative variation inherently implied by these growth forms. Distilling coral morphology into geometric variables that capture shape variation will allow for better tests of the mechanisms that govern coral biology, ecology and ecosystem services such as reef building and provision of habitat.

## INTRODUCTION

The shape and size of organisms determines how they interact with the physical environment and with other organisms (Denny 1993; Vogel 1996). This is especially true for sessile colonial organisms, where variation in morphology has been linked to a range of biological and ecological processes (Jackson 1977, 1979). For example, growing upwards from the benthos avoids benthic competition, whilst growing laterally avoids whole colony mortality by spreading risk (Jackson 1979). Despite the fundamental importance of a colony’s morphology, there is no general framework for capturing morphological variation. Instead, scientists tend to lump individuals and species into discrete growth form categories (corals: (Veron 2000); bryozoans: (Bishop 1989); bacteria: (Shapiro 1995)) or use continuous metrics that cannot partition the effect of size and shape (e.g., surface area to volume ratio; Naumann et al. 2009). Developing a quantitative framework is challenging for colonial organisms because they have geometrically complex forms, high intraspecific and interspecific variation in shape, and lack readily identifiable landmarks for comparative analysis. However, new technologies such as CT and laser scanning make it possible to accurately capture the diversity of shapes exhibited by colonial organisms (Lavy et al. 2015; House et al. 2018). Here, using reef corals as a study system, we develop a morphological schema using quantitative, three-dimensional shape variables.

Scleractinian corals are a prime example of colonial organisms whose morphology directly dictates life history strategies (Jackson 1979), demographic rates (Dornelas et al. 2017; Álvarez-Noriega et al. 2016; Madin et al. 2014) and provisioning of habitat for other taxa (Bell and Galzin 1984; Richardson et al. 2017; Graham and Nash 2013). The aragonite skeleton that scleractinian corals secrete as they grow provides support and shape, however most of the live biomass is associated with the surface (Hoegh-Guldberg 1988; Johannes and Wiebe 1970). These characteristics have consequences for vital processes such as growth and survival, where higher surface area to volume ratios allow more biomass per unit investment in skeleton, but may increase the risk of partial colony mortality (Lirman 2000), dislodgement (Madin 2005), and susceptibility to thermal bleaching (Baird and Marshall 2002). Meanwhile, coral structures provide habitat for many taxa and can act as predator refuge for both adult and juvenile fishes (Kerry and Bellwood 2012a; Friedlander and Parrish 1998). The diverse and complex morphologies produced by corals also changes the local environmental conditions, such as light (Sheppard 1981) and water flow (Hench and Rosman 2013), to create a range of microhabitats and niches. At the habitat scale, variation in the morphology of each colony in an assemblage contributes to the overall complexity of the habitat while alive or via the skeleton left behind after it has died (Richardson, Graham, and Hoey 2017), with variation in structural complexity linked to ecosystem properties, such as microhabitat availability (Graham and Nash 2013) which in turn influences fish body size distributions and adult fish assemblage structure (Almany 2004; Nash et al. 2014). Furthermore, the aragonite structures built by the corals are the main component of the reef superstructure, where variation in colony shape influences the persistence of colony skeleton following mortality and reef matrix building and infilling processes (Glynn and Manzello 2015; Rasser and Riegl 2002). Yet, despite the importance of morphology for the functioning of both the corals themselves and coral reef ecosystems, quantitative studies of coral morphology are sparse, presumably because of difficulties in measuring and dealing with the geometric complexity of coral forms.

Scleractinian corals exhibit high levels of variation in morphology within and among taxa. They vary from simple shapes, such as encrusting or hemispherical colonies, to tree-like branching shapes. There are also varying degrees of morphological plasticity within species driven by interactions with local environmental conditions (Foster 1979). Additional phenomena such as partial mortality (Meesters, Wesseling, and Bak 1996), colony fragmentation (Karlson 1986), and indeterminate growth (Sebens 1987) add to the complexity and observed variation in morphology from colony to colony, even within species and conspecifics. Corals need access to free-flowing water for filter feeding, and light for photosynthesis, both of which are linked to morphological variation (Hoogenboom, Connolly, and Anthony 2008; Kaandorp et al. 1996). Additionally, competition for space results in many colonies growing up from the substrate to increase standing biomass without needing to continuously colonize new substrate (Jackson 1977). However, many sessile colonial organisms within marine environments are subjected to hydrodynamic forces that can dislodge entire colonies if they grow too far away from the substrate, restricting the range of available morphologies (Koehl 1999). Taken together, coral colonies exhibit multiple morphological trade-offs that result in the vast array of observed variation in morphology (Kaandorp et al. 1996; Chappell 1980).

Scleractinian corals are typically categorised into growth forms based on coarse morphological similarities. Growth forms are useful for species identification and monitoring bulk change in assemblage structure, but do not adequately capture geometric complexity or intraspecific variation in shape. Phenotypic plasticity is common among coral species, where the same species can exhibit different growth forms in different environments (Veron 2002). Despite these limitations, growth form is still a useful metric because life processes and functions tend to differ significantly among categories. For example, growth form is a good predictor of competitive ability (Connell et al. 2004; Hoogenboom, Connolly, and Anthony 2008) and zonation patterns (Chappell 1980; Done 2011). Recent work has highlighted the link between growth form and size with demographic rates including fecundity, growth, and background mortality (Álvarez-Noriega et al. 2016; Dornelas et al. 2017; Madin et al. 2014). However, while these results highlight differences between growth forms across a range of processes, they are unable to directly assess process-based hypotheses for the observed differences, nor can the results be generalised to other growth forms or taxa with similar morphological adaptations but different overall morphology (e.g., sponges, hydrozoans, algae, plants, etc). As such, recent studies have begun to explore techniques for quantifying and comparing the three-dimensional shape of corals (Lavy et al. 2015; Reichert et al. 2017; Bythell, Pan, and Lee 2001; House et al. 2018). We build on this work to develop a quantitative schema for coral morphology via variables that capture shape variation.

Previous work has shown that size-dependent relationships vary between growth forms (Madin et al. 2014; Dornelas et al. 2017), suggesting that variation in shape is important. Quantitative variables that attempt to explain these differences should therefore aim to partition the effects of shape and size separately. However, surface area to volume ratio changes with both size and shape: two colonies with the same shape but different volumes will have different surface area to volume ratios, and two colonies with the same volume and different shapes can also have different surface area to volume ratios. Surface rugosity, on the other hand, is size-invariant. However, it only captures the upper surfaces of colonies, potentially missing key aspects of shape variation within or underneath colonies. There has also been a lack of comprehensive studies that measure morphology across a broad range of colony shapes and sizes and along multiple axes of variation simultaneously. By measuring multiple, size-independent variables across a wide range of morphological variation, trade-offs and broader patterns become clearer. These variables should then be useful as functional traits, defined as a measurable property of an organism linked to performance (McGill et al. 2006; Violle et al. 2007), that can be used to provide mechanistic explanations for variation in colony biology and ecology.

Variables that can capture how a colony is spatially distributed in the environment should capture functionally relevant axes of morphological variation. For example, variables that capture how compact the volume of a colony is may act as a good indicator of “branchiness”, how sturdy a colony is, or act as a proxy for a size-independent surface area to volume ratio, capturing a continuous gradient from massive to arborescent colonies. Variation in compactness may therefore capture a continuous gradient that covaries with processes such as growth rates, fragmentation, and habitat provision (Gladfelter, Monahan, and Gladfelter 1978; Lirman 2000; Alvarez-Filip, Gill, and Dulvy 2011). Another example is how top heavy a colony is, which would capture how surface area or volume is distributed away from the substrate, capturing a continuous gradient from low-lying encrusting to tabular colonies. This axis can be expected to covary with processes such as whole colony dislodgment, benthic competition strategy, and microhabitat diversity (Madin et al. 2014; Jackson 1979; Kerry and Bellwood 2012b). Yet another axis of variation is how the surface area of a colony is distributed, which should capture a gradient from smoother, less convoluted surfaces to colonies with highly complex and convoluted surface area distribution. Variation in surface complexity may capture a functional trade off axis between biomass packing (e.g. having more biomass for a given area of space) and decreased intra-colony competition for resources (e.g. increased light per unit biomass when surface area is spread out) (Wangpraseurt et al. 2012; Hoogenboom, Connolly, and Anthony 2008). By measuring these axes of morphological variation, we can place coral colonies along multiple functional axes, moving from a subjective, categorical framework towards and quantitative, functional trait-based one.

Morphology is important for corals and the ecosystems they build, but a comprehensive suite of quantitative traits for developing explanatory and generalised models for these processes have yet to be formalised. The aim of this study was to develop a set of generalised morphological variables that capture biologically relevant axes of variation in corals. To achieve this aim, we first derived six morphological variables and measured them across a broad diversity of colony shapes and a wide range of sizes via high-resolution 3D laser scanning. We then asked: (i) how do corals occupy continuous morphological space? (ii) where do growth form classifications sit within this space? (iii) how does the shape of growth forms covary with size? And, (iv) do continuous variables capture the subjective information encoded in growth forms? We show that the variables outlined in this study can place growth forms on three continuous axes of variation and provide a more precise, mechanistic toolkit for ongoing research.

## MATERIALS AND METHODS

### Data collection

Colony skeletons from coral collections at the Natural History Museum in London (UK), the Bell Pettigrew Museum at the University of St Andrews (UK), and the Museum of Tropical Queensland (Australia) were scanned using an optical laser scanner (EXAScan, Creaform.inc) to digitize their threedimensional morphology. Colonies were selected for scanning based on being mostly intact and capturing a broad range of shapes and sizes across a diversity of traditional coral growth form classifications (arborescent, corymbose, digitate, laminar, massive, sub-massive, or tabular; (Veron 2000)).

Scanning was conducted with a standard resolution of 0.5 mm^2^ but needed to be decreased to 1 mm^2^ for several large or complex colonies due to computational constraints. All colony scans were orientated with the z-axis aligned with the colony’s likely upward orientation when on the reef. Each scan consisted of a digital 3D mesh that was comprised of a single contiguous surface of connected triangles. Meshes were rejected if the final mesh deviated in shape from the actual specimen due to issues associated with interpolating missing scan data. The final dataset consisted of 152 meshes. To test the precision and accuracy of the laser scanner, 20 colonies previously scanned using a medical CT scanner (House et al. 2016) were rescanned using the laser scanning protocol and morphological measurements were compared (Fig. S1).

### Size and shape variables

For each colony mesh, we calculated two size variables, volume and surface area, and six shape variables: sphericity, convexity, packing, fractal dimension, and the second moments of area and volume.

Sphericity *S* is a size invariant measure of the compactness of an object’s volume (Wadell 1935). It is calculated as the ratio of the surface area of a sphere with the same volume as the object and the surface area of the object. Sphericity is bounded by zero (i.e., a theoretical shape that is entirely non-compact, like a plane) and one (i.e., a perfect sphere) and is size independent. Because sphericity is a ratio between zero and one, but never exactly zero or one, it was logit-transformed for analyses.

Convexity *C* is a size invariant measure of the degree to which there is space between different parts of an object (Zunic and Rosin 2004). Convexity is calculated as the volume of an object divided by the volume of its convex hull, where the convex hull is the shape formed by the smallest possible boundary that has no concave areas around an object (Barber, Dobkin, and Huhdanpaa 1996). Like sphericity, convexity is bounded by zero (i.e. a theoretical shape that has no volume but some convex volume) and one (i.e. a shape that is entirely convex) and was logit-transformed for analyses.

Another form of convexity (which we call packing *P* for clarity), is a size invariant ratio of how much of an object’s surface area is situated internally versus externally in relation to its immediate environment (Zunic and Rosin 2004). It is calculated as the surface area of an object divided by the surface area of its convex hull. A packing value above one indicates that surface area is packed within the volume it occupies (i.e., it is more inverted). Values below one indicate that surface area is more spread out over the volume it occupies (i.e., it is more everted). Convex shapes have a packing equal to one, as the surface area is neither internally nor externally distributed, and objects with an equivalent amount of “inverted” and “everted” surface also have a packing equal to one. As packing is a proportion that can go above or below 1, it was log^10^ transformed for analyses.

Fractal dimension *D* captures how surface area fills space and is an estimate of spatial complexity. We calculated fractal dimension using the “cube counting” algorithm, a 3D analogue of the well-known box counting method (Sarkar and Chaudhuri 1994). Fractal dimension is bounded between two (a plane) and three (a theoretical 2D surface that is completely volume filling) and is size invariant. Fractal dimension is calculated as the slope of log^10^(*N*)~log^10^(*C*), where *N* is the total number of cubes that contain any surface of the object and *C* is the number of equal sized cubes in the 3D cube array.

The previous four shape variables are all rotationally and size invariant (i.e., the orientation or size of colony meshes has no bearing on the resulting value). However, a distinguishing feature of shape in corals is how volume and surface area are distributed vertically above the substrate. For instance, a tabular coral colony has volume and surface area distributed further away from the substrate (i.e., “top heavy”) than a hemispherical colony that is “bottom heavy”. To capture this feature, we used second moments of volume (V_VOL_) and surface area (V_AREA_), which are the sums of the products of volume and area, respectively, with their distance from the colony attachment point. Both variables were log^10^ transformed for analysis. To ensure size invariance, colony meshes were converted to a standard volume of 1mm^3^ before these variables were calculated.

Size variables, surface area and volume, were calculated on the original sized meshes. Surface area was calculated using the total surface area of the colony, minus any surface area that would be attached to the substrate.

### Analysis

We used principal components analysis (PCA) to visualize the morphospace, how growth forms occupied this space, and to identify which shape variables explained most of the variation in colony shape (via the ‘prcomp’ function in R (R Core Team 2015)). Variables were standardized with a mean of zero and unit variance to reduce the influence of variable scale on the projection. For each principal component, variables were highlighted as important for a given component based on whether their loadings exceeded the null contribution value of 16.6% (100% divided by six variables). Pair-wise plots of raw data with Pearson’s correlations were used to identify how variables covary and which variables were highly collinear both within a given component and between the variables overall.

To test whether shape remained constant with colony size we used a linear regression approach, with shape variable as the response, and growth form, volume and their interaction as predictors (using the ‘lm’ function in R). Each shape variable within each growth form were deemed to remain constant with size if zero was within the 95% confidence intervals for the slope estimate.

To infer whether the morphological variables captured a broad proportion of the subjective variation encoded in growth forms, we first added 95% confidence ellipses for each growth form to the PCA to visualize how growth forms occupied continuous shape space. We then used multinomial regression to see how well the shape variables could predict the correct growth form (via the ‘multinom’ function from the R package ‘nnet’ (Venables and Ripley 2002)). We built the initial model based on a set of variables that captured the main axes of variation in shape and had low covariance to minimise redundancy of information. No interactions between shape variables were included. Additionally, volume was not included as a main effect as size is not a determining characteristic for growth form. However, volume was included as an interaction effect with shape variables that were shown to vary as a function of volume. Finally, we used this model as the basis for a leave-one-out assessment of predictive accuracy, where a model that omitted an observation was fitted, and the omitted observation data used to generate probabilities of that observation belonging to any one growth form classification, as well as assigning a single class. This was repeated for every observation sequentially and the predicted probabilities and classes were pooled, with the probabilities used for a visual assessment of model performance and the classes used for the generation of a confusion matrix to assess classifier (in this case the multinomial model) performance. The predicted class dataset was assessed via kappa values and balanced accuracy estimates to assess classifier accuracy (Cohen 1960). The overall model was assessed for goodness of fit via McFaddens pseudo-R^2^.

## RESULTS

### Corals in continuous shape space

88% of the observed variation in shape was captured by the first two principal component (PC) axes (Fig. 1). The first PC axis captured 60% of the variation across the six-dimensional shape space and was comprised of sphericity, convexity and the second moment of area (V_AREA_). All three variables had contribution values above 16% with sphericity and V_AREA_ having joint highest at 25% and convexity at 22%, suggesting they were all important for explaining the variation along PC1. All three variables were highly correlated (Fig. 2), where sphericity and convexity were positively correlated, with both being negatively correlated with V_AREA_. Of these three variables, convexity was selected for the predictive model as it had the weakest correlation with the other variables, therefore minimizing redundancy. The second PC axis captured 27.5% of the variation and was comprised of packing, fractal dimension and the second moment of volume (V_VOL_). Of these variables, packing had the highest contribution at 35%, followed by fractal dimension (32%), and V_VOL_ (24%). Packing and fractal dimension were highly correlated with each other (Fig. 2); however, both were uncorrelated with V_VOL_. Of these variables, both V_VOL_ and packing were selected for the predictive model; V_VOL_ because it was uncorrelated with the other two and packing as it was slightly less correlated with the other variables compared to fractal dimension.

**Fig 1.**
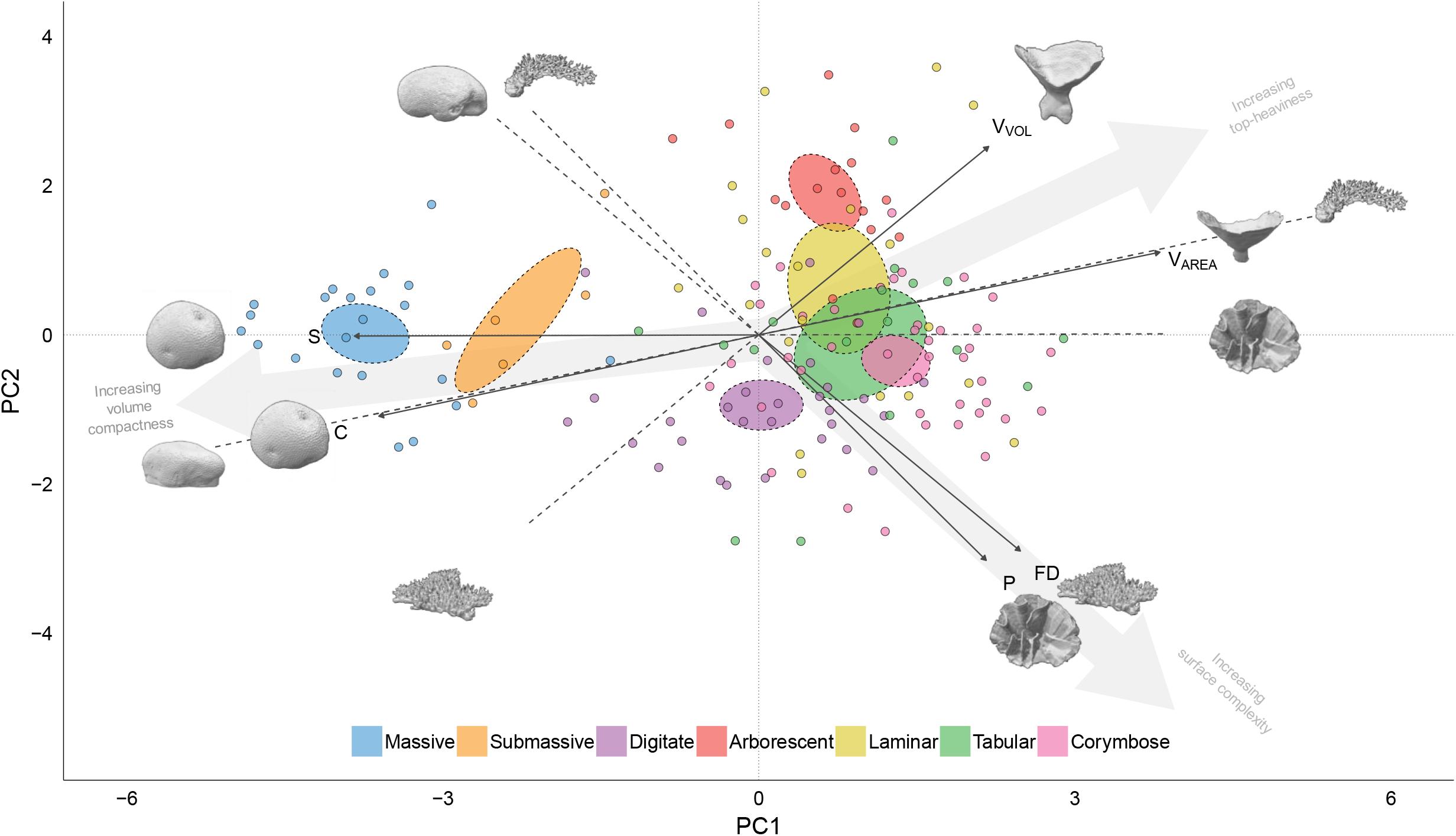
Projection of 152 coral colonies in two dimensions by the 1st and 2nd principal component (PC) axes of six-dimensional shape space. Points colored by growth form classification with 95% confidence ellipses around the group means. Arrows indicate the loading and direction of each shape variable; V_VOL_ = 2nd moment of volume (mm^4^), V_AREA_ = 2nd moment of area (mm^3^), FD = fractal dimension, P = packing, C = convexity, S = sphericity. The first principal component broadly captures variation in skeletal volume compactness (S, C & V_AREA_). The second principal component captures a trade-off between surface area complexity (FD & P) and the distribution of volume vertically in the water column (V_VOL_). Images are of the coral specimens that occupy the extremes of each shape variable in the dataset, with some specimens occupying the extreme ends of multiple variables.

**Fig 2.**
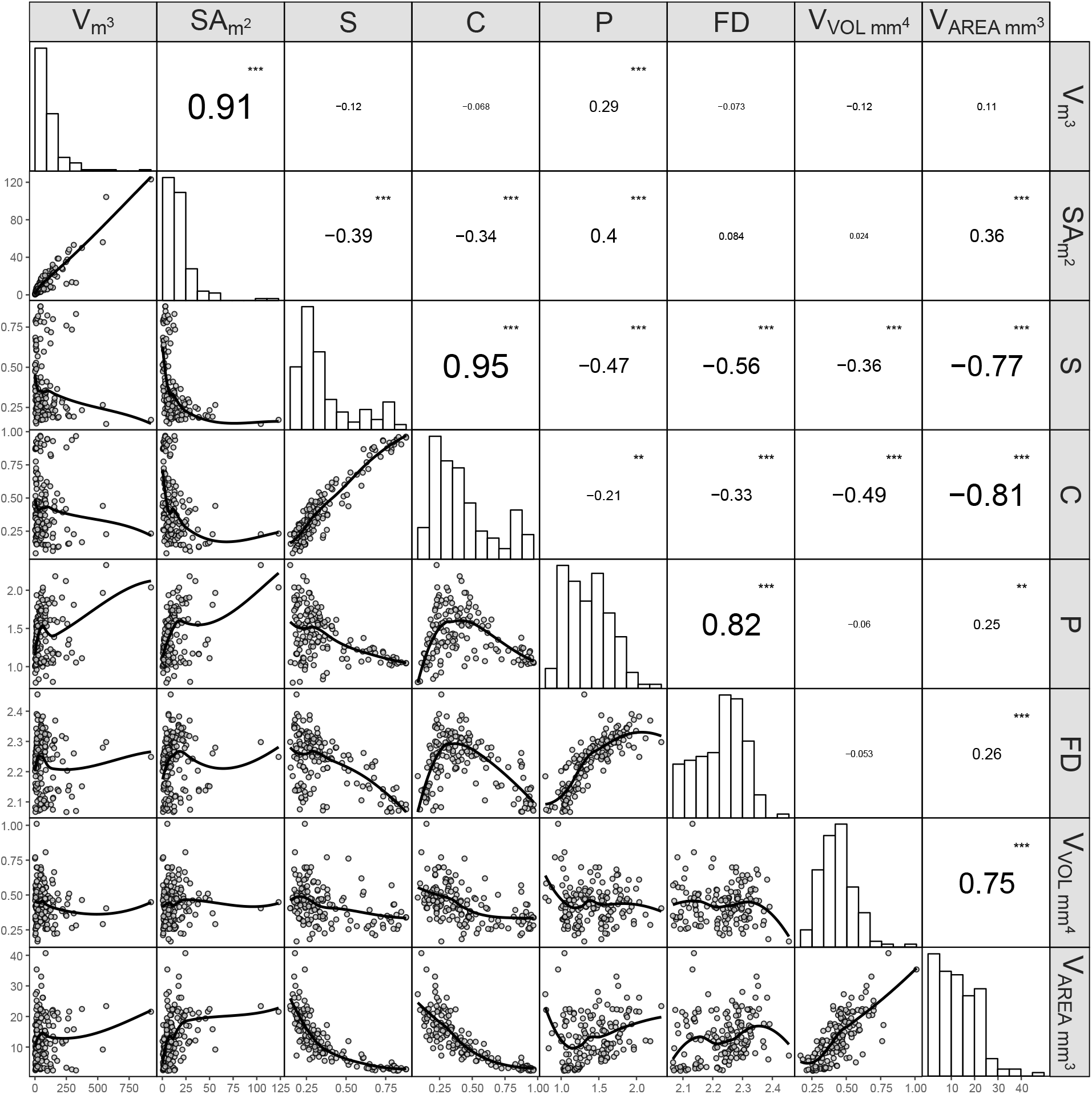
Pair-plot of six shape and two size variables used in the study. V = volume, SA = surface area, S = sphericity, C = convexity, P = packing, FD = fractal dimension, V_VOL_ = 2nd moment of volume (mm^4^), V_AREA_ = 2nd moment of area (mm^3^). Bottom triangle panels: Scatter plots of each variable pair with loess smoother line, n = 152. Diagonal panels: density plots of each variable, Upper triangle panels: Pearson’s correlations for each variable pair with significance scores (*** = p < 0.001, ** = p < 0.01).

Coral colony shape is constrained by compactness. When sphericity and convexity are high, there was less variation in complexity (captured by packing and fractal dimension), with a similar but less pronounced effect on V_VOL_. (Fig. 2). Additionally, there was a strong, exponential decrease in V_AREA_ as a function of these two variables. Sphericity had the highest correlation scores with the other shape variables. In the PCA projection we also observed this constraining effect, where the spread of points along PC2 is markedly restricted in extent and density at lower PC1 scores (i.e. higher sphericity and convexity) (Fig. 1). As sphericity and convexity decreased however, the extent of occupied shape space along PC2 increased.

### Growth forms in continuous shape space

There were two apparent gradients that captured how growth forms were distributed in continuous shape space (Fig. 1). The first was along PC1 where the massive and sub-massive growth forms were isolated from the branching growth forms. The second was along PC2 within the branching group, with the digitate, corymbose, tabular, laminar and arborescent growth forms distributed roughly in that order. The mean position for a given growth form overall was generally constrained within shape space, however at the colony level, many growth forms were found occupying the same area which was partially explained by variation in shape as a function of size.

### Changes in colony shape with size

While there were no significant correlations between any shape variable and size (represented as colony volume) across all the observed values together except for packing (Fig. 2), the shape of a colony did change as a function of size within some growth form and shape variable combinations (Fig. 3). Sphericity values decreased with size in the digitate and laminar growth forms. All other growth forms maintained constant sphericity across their observed size range except the tabular and arborescent group which appeared to decrease marginally as volume increased despite their slope estimate confidence intervals overlapping with zero. Packing increased with size fastest in the arborescent group, followed by the corymbose, laminar, digitate and tabular growth forms, with the massive and sub-massive colonies remaining constant. The massive group decreased in fractal dimension with size and the corymbose colonies increased with size. All the other variables remained constant as a function of volume based on the 95% confidence intervals. However, there was some evidence to suggest that both V_VOL_ and V_AREA_ may scale with size in some growth forms.

**Fig 3.**
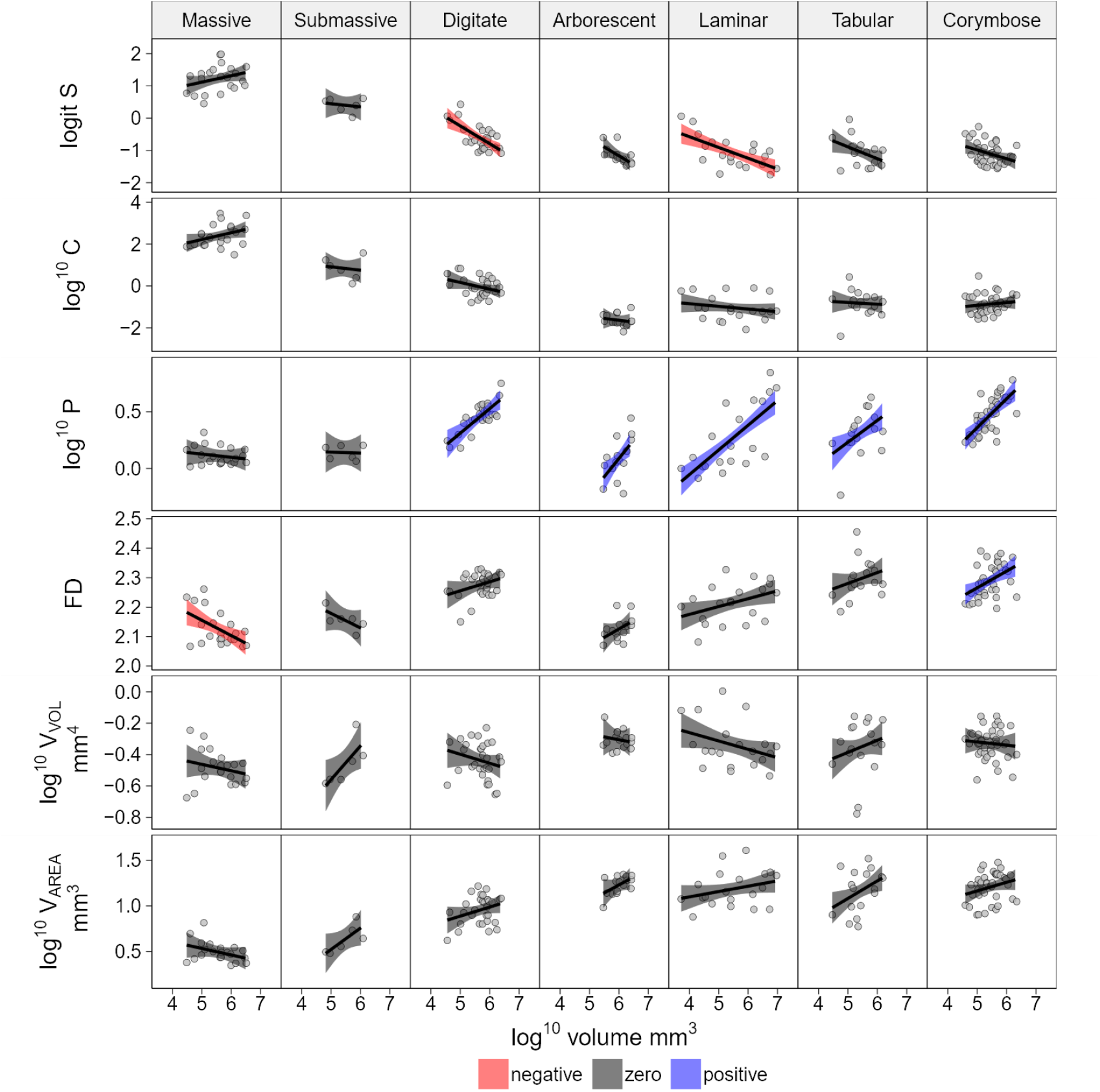
Size by shape variable plot for 152 coral colonies faceted by growth form highlighting changes in shape as a function of colony volume. Panel order from left to right based on average PC1 values for each growth form. Lines represent linear regression lines with 95% confidence intervals colored based on whether the 95% confidence intervals for slope estimates overlapped with zero. S = Sphericity, C = Convexity, P = Packing, FD = Fractal dimension, V_VOL_ = 2^nd^ moment of volume, V_AREA_ = 2^nd^ moment of surface area.

### Capturing qualitative growth forms using quantitative variables

Growth form was correctly predicted by four shape variables in conjunction with volume (Fig. 4). The model included convexity, packing, V_VOL_ and fractal dimension, with interaction terms between volume and both packing and fractal dimension. The final model explained 74% of the deviance (McFadden’s pseudo R^2^ of 0.62, d.f = 42). Overall the model predicted growth forms with a high degree of accuracy (kappa = 0.66). The growth forms in order of highest to lowest balanced accuracy were; massive (95.1%, n = 22), arborescent (92.6%, n = 16), sub-massive (91.3%, n = 6), digitate (81%, n = 30), corymbose (80.7%, n = 41), tabular (77.2%, n = 17) and laminar (70.2%, n= 20).

**Fig 4.**
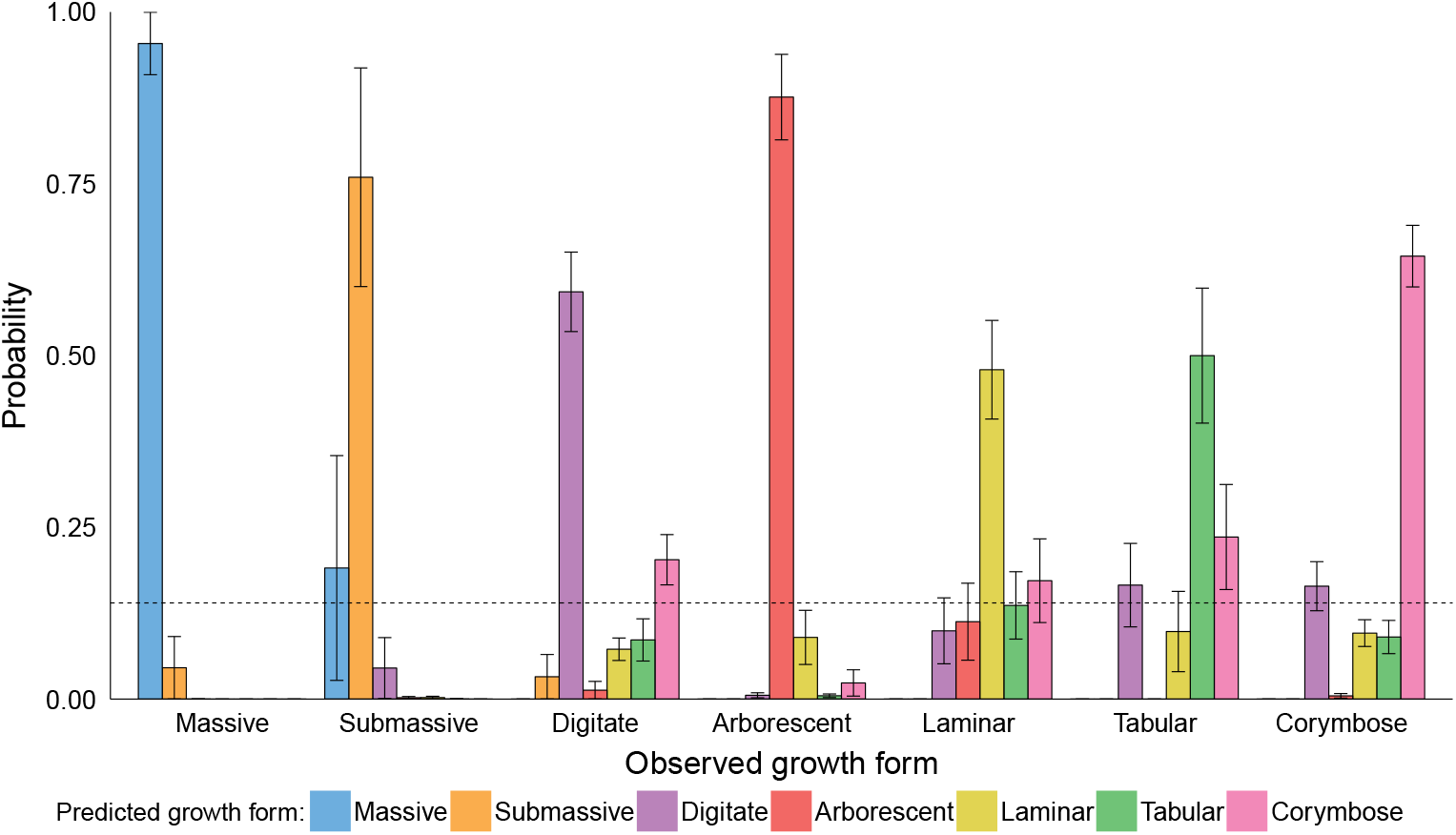
Observed growth form by predicted growth form probabilities for seven coral growth forms based on a multinomial regression using continuous shape variables. Data generated via a leave-one-out approach, where each observation was left out of the initial model and classification probabilities generated for the missing observation, repeated for each observation in the dataset. Colored bars represent average probability of being classified as a given growth form with standard errors. Horizontal dashed line represents the unbalanced expected probability if all classes were randomly assigned (100%/7 possible classes) and was used to determine which incorrectly predicted classes were significantly misclassified for each growth form. n = 152.

Fractal dimension was added to the model as the three shape variables alone were unable to distinguish between tabular and laminar growth forms, despite the balanced accuracy of all other groups being above 79%. The probability of the correct class in the final model was the highest for all growth forms and was distinct from all other potential classes (Fig. 4). The massive group had the highest mean probability (0.95 ± 0.04), with the laminar and tabular having the lowest (0.48 ± 0.07 and 0.50 ± 0.1, respectively).

## DISCUSSION

We developed six quantitative shape variables and showed that variation in volume compactness, surface complexity and top-heaviness explain much of the variation in coral shape. All shape variables were size invariant, and so the observed changes in some variables as a function of colony size demonstrates specific aspects of colony morphology that are more likely to change as individuals grow, which also leads to colonies that straddle traditional growth form classifications. We found that four morphological variables can predict traditional growth form classifications with accuracies ranging from 70 to 95%, therefore demonstrating that these variables capture a broad amount of the subjective variation encoded in traditional growth form classifications. Given that morphology has been shown to be an important predictor of biological and ecological function in corals, and relates to their role as ecosystem engineers, the ability of quantitative shape variables to delineate growth form also suggests an ability to capture continuous functional axes. our approach is suitable for placing coral colony morphology along continuous axes of functional variation without relying on homologous structures or landmarks, providing a set of morphological traits to test generalised and mechanistic hypotheses for biological and ecological processes.

Corals occupy continuous shape space along three main axes of variation. Variation in volume compactness captures a gradient from non-branching to highly-branching colonies. However, compactness constrains surface complexity and top-heaviness, where colonies with higher levels of compactness tend to be smooth and bottom heavy. Furthermore, each of the three main axes of variation can provide general, mechanistic explanations for biological processes. For example, volume compactness, capturing the gradient from massive to more complex forms, may be a suitable trait for explaining why more massive morphologies have less variable and slower overall growth (Dornelas et al. 2017). Similarly, variation in top-heaviness, capturing a gradient from lower lying to tabular colonies, may be a trait that can test ideas related to benthic competition strategies (e.g. lateral benthic expansion vs indirect shading and competitive escape) (Jackson 1979). Variation in surface complexity is related to competition and resource use, where colonies with their surfaces distributed in a complex way have less resources (e.g. light, nutrients) per unit surface area but can have more polyps packed within a given space (Wangpraseurt et al. 2012). These hypotheses are based on organism performance, but others can be formulated across a range of scales. Examples include low compactness related to more space for juvenile fishes (Alvarez-Filip, Gill, and Dulvy 2011), high compactness increasing reef framework building (Rasser and Riegl 2002), high surface complexity increasing larval recruitment (Hata et al. 2017), and niche diversification being increased by top-heavy, tabular colonies (Kerry and Bellwood 2015). The ability to formulate and test these types of process-based hypotheses offers a direct approach for linking form to function not possible using growth forms or clouded by metrics that conflate shape and size.

More broadly, the interaction between surface area and volume is important across taxa (Folkman & Hochberg, 1973; Jackson, 1979; Tilkens, Wall-Scheffler, Weaver, & Steudel-Numbers, 2007). In such cases, sphericity should be used instead of surface area to volume ratios as sphericity captures surface area to volume ratios in a size independent manner, avoiding conflating shape and size effects due to the isometric scaling of surface area and volume, and allowing for decoupled exploration of how shape changes with size. For example, microbes and bacteria have a variety of morphologies that modulate a variety of processes such as nutrient uptake, motility, and predator avoidance, and both size and shape play a key role (Young, 2007). The cost of switching to sphericity is nil given that both sphericity and surface area to volume ratio require the same data to calculate.

Growth forms typically occupy specific areas of continuous shape space, but the large amount of overlap between them suggests that growth forms are less distinct than their discrete nature implies. Growth forms were also not distributed along a single trajectory of morphological variation which highlights that morphological variation between growth forms occurs along multiple axes. Therefore, a single ordinal classification of categories, for example from most to least “complex”, would be misguided. Overall, the semi-distinct, semi-overlaying distribution and variation in growth forms at once confirms that growth forms work as morphologically distinct classifications to some degree but at the same time suggests that they are less definite and defined than implied by the nature of assigning a single category. This potentially unsatisfactory statement is made more palatable once morphological plasticity and the observed changes in shape as a function of size within growth form are taken into account: not every corymbose colony looks alike in the same way that each member of a wildebeest herd does, and a tabular colony only looks truly tabular after an initial period of growth up and out from the benthos. As such, the size distribution of colonies may have implications for the shape distributions of colonies within a growth form. Within growth form changes in life history traits with size have been highlighted previously (Álvarez-Noriega et al. 2016; Dornelas et al. 2017; Madin et al. 2014), which may be partially explained by ontogenetic changes in morphology and morphology-related processes. While variation in processes between growth forms act as an indicator of a morphology-process link, the incomplete and overly definite nature of growth form categories are unsuitable for establishing causal links.

While this study included a wide range of growth forms and sizes, there are unobserved sources of variation that may fill out or stretch the boundaries of the observed shape space if added. Encrusting colonies, which extend laterally over the surrounding substrate, were not included due to the difficult nature of obtaining whole colony specimens and the fact that the three-dimensional shape of an encrusting colony is contingent on the local substrate it encrusts, although *in situ* measurements of encrusting forms should be possible via photogrammetry techniques. Columnar colonies are also absent due to a lack of intact specimens in the museum collections. The less populated space between the massive and sub-massive growth forms and the remaining growth forms would likely be occupied by columnar colonies given their semi-sturdy, semi-branching shape. While the range in colony size in the study varied over three orders of magnitude, including both smaller and larger colonies in the dataset may also further fill in and expand the observed shape space.

Both the approach and results of our study have applications for relating morphology to process in other taxa. Because the variables used in this study require no taxon-specific information to calculate (e.g. landmarks) they can be used to measure and compare morphological variation across any organism where a suitable 3D representation is available. Other colonial organisms such as sponges, soft corals, gorgonians and macroalgae are similar in their range of geometric complexity and are exposed to similar conditions, which suggests that similar trade-off axes should exist in these taxa. Measuring complex colony shapes across taxonomic groups allows for empirical testing of the theoretical work on morphological strategies laid out by Jackson (1979). In the terrestrial realm, there is a large body of work on the functional ecology of plants which partially overlaps with corals given that both groups are sessile, able to experience partial mortality, and have a photosynthetic component. Going a step further, it should also be possible to compare the morphology across a broad spectrum of organisms, from bacteria to blue whales, to uncover more universal drivers of morphological adaptations via the variables outlined in this study.

This study provides a comprehensive set of traits that partition shared and unique variation between growth forms and highlight size-dependent changes in shape within growth forms. These traits have strong theoretical links to many processes important for both corals and their roles as ecosystem engineers and allow for mechanistic and generalised explanations of phenomena to be established. This study provides an empirical toolkit and theoretical backbone for future reef research that is timely given the ongoing work on three-dimensional metrics and methods, and the need to establish a broader understanding of how morphology maps to function across scales.

## DATA ACCESSIBILITY

Data and r scripts used in this manuscript will be uploaded to figshare.

## AUTHORS’ CONTRIBUTIONS

All authors designed the study. K.Z. collected and analysed the data, created figures and wrote the first draft. All authors contributed substantially to revisions.

## ACKNOWLEDGEMENTS

We would like to acknowledge Francois Leclerc and CREAFORM Inc. for providing equipment and technical support for the data collection. Luc Bidaut and Jenny House for CT scan data for calibration. Miranda Lowe and the Natural History Museum of London, Tom Bridge and the Museum of Tropical Queensland, and the Bell Pettigrew museum for facilitating specimen access and data collection. Kristin Precoda, Rachael Woods and Damaris Torrez-Pulliza for proofreading and feedback throughout.

